# Foot shock facilitates reward seeking in an experience-dependent manner

**DOI:** 10.1101/2020.06.04.134783

**Authors:** JA Strickland, AD Dileo, M Moaddab, MH Ray, RA Walker, KW Wright, MA McDannald

## Abstract

Animals organize reward seeking around aversive events. An abundance of research shows that foot shock, as well as a shock-associated cue, can elicit freezing and suppress reward seeking. Yet, there is evidence that experience can flip the effect of foot shock to facilitate reward seeking. Here we examine cue suppression, foot shock suppression and foot shock facilitation of reward seeking in a single behavioural setting. Male Long Evans rats received fear discrimination consisting of danger, uncertainty and safety cues. Discrimination took place over a baseline of rewarded nose poking. With limited experience, all cues and foot shock strongly suppressed reward seeking. With continued experience, suppression became specific to shock-associated cues and foot shock facilitated reward seeking. Our results provide a means of assessing positive properties of foot shock, and may provide insight into maladaptive behavior around aversive events.

## Introduction

Animals must navigate a perilous world to secure essential rewards. Studies of associative learning and defensive behaviour reveal that animals are adept at organizing reward seeking around aversive events. Even more, this organization changes as a function of experience. Picture a mildly hungry rat responding on a lever for food. Presenting an innocuous cue will produce little change in the rat’s reward seeking behaviour. A foundational report by Estes and Skinner (1941) demonstrated associative learning of a cue paired with foot shock endows the shock predictive cue with aversive properties^1^. A shock-associated cue will sharply reduce reward seeking. Historically termed a conditioned emotional response^2–5^, now commonly termed conditioned suppression, the ability of a shock-associated cue to suppress reward seeking has been observed in many settings and laboratories^6–16^.

Like their predictive cues, aversive events alter behaviour. Picture a rat exploring a novel environment when foot shock is unexpectedly delivered. Foot shock delivery will elicit a brief, undirected activity burst, followed by freezing^17–20^. While theoretical accounts vary^21–23^, the behavioural phenomenon of post shock freezing is widely observed^24–28^. Foot shock can also suppress reward seeking^29^, and the organization of reward seeking around foot shock changes with experience^30^. In their original demonstration, Estes and Skinner noted a compensatory increase in responding following shock delivery, though this was perhaps a return to baseline. LaBarbera and Caul (1976) later demonstrated that foot shock can facilitate reward seeking^19^. Even more, the magnitude of facilitation increased with foot shock experience. These findings indicate that reward seeking around an aversive event may change dramatically with experience.

Studies of cue suppression of reward seeking, post shock freezing, and post shock facilitation of reward seeking are typically carried out in isolation. These studies have utilized different dependent measures of behaviour: ratios derived from rates of reward seeking, freezing, and instrumental response rates. By necessity, these studies differed in basic experimental details: foot shock intensity, foot shock duration, cue type, cue length, etc. Our laboratory has developed a conditioned suppression procedure in which distinct auditory cues predict unique foot shock probabilities: danger (*p*=1.00), uncertainty (*p*=0.25) and safety (*p*=0.00). Cues are presented over a baseline of rewarded nose poking, and poke-reward contingencies are independent from cue-shock contingencies. Using this procedure, we have observed a consistent, experience-dependent pattern of discrimination. Initial suppression of nose poking to all cues gives way to graded responding that reflects shock probability: danger > uncertainty > safety^31–36^.

The goal of the current study is to examine cue suppression, post shock suppression and post shock facilitation of reward seeking in a single behavioural setting. To fairly compare each mechanism, we use a common dependent measure: nose poke rate. Nose poke rate is particularly suitable because it is objective and permits second-by-second analysis of responding. By examining nose poke rate in discrete time periods across discrimination, we are able to reveal how experience shapes reward seeking around aversive shock and shock-associated cues.

## Materials and Methods

### Subjects

Subjects were 123 male Long Evans rats pooled from 9 different studies (n = 15, 17, 13, 11, 21, 7, 21, 8, 10). 110 were obtained from Charles River (26 shipped ~21 day olds, 84 ~50 days old) and 13 were born in the laboratory. Those born in the ACF were housed with mothers until postnatal day 21 when they were weaned and all single housed. All were maintained on a 12-hour light-dark cycle (lights on 0600–1800) and were aged approximately 74 - 84 days at the start of fear discrimination, throughout which they were maintained at 85% of their free-feeding body weight and received water ad libitum. Eighty-four subjects received sham lesions prior to fear discrimination in the following brain regions: lateral habenula (n=17), orbitofrontal cortex (n=21)^33^, ventral pallidum (n=7), nucleus accumbens core (n=21)^36^, retrorubral field (n=14), and dorsal raphe (n=4). Twenty-four of the rats not given sham lesions received additional handling in adolescence.The remaining fifteen rats had access to a second water bottle during adolescence. All protocols were approved by the Boston College Animal Care and Use Committee, and all experiments were carried out in accordance with the NIH guidelines regarding the care and use of rats for experimental procedures.

### Apparatus

The apparatus for fear discrimination consisted of eighteen individual sound-attenuated enclosures that each housed a behaviour chamber with aluminum front and back walls, clear acrylic sides and top, and a metal grid floor. Each grid floor bar was electrically connected to an aversive shock generator (Med Associates, St. Albans, VT). A single food cup and central nose poke opening equipped with infrared photocells were present on one wall. Auditory stimuli were presented through two speakers mounted on the ceiling of each enclosure.

### Nose poke acquisition

All rats were first provided pellets (Bio-Serv, Flemington, NJ) for two days in the home cage. Rats were then shaped to nose poke for these pellets in the experimental chamber. During the first session, the nose poke port was removed, and rats were issued one pellet every 60 seconds for 30 minutes. In the next session, the port was reinserted, and poking was reinforced on a fixed ratio 1 schedule in which one nose poke yielded one pellet until they reached ~50 nose pokes. Nose poking was then reinforced on a variable interval 30-second (VI-30) schedule for one session, then a VI-60 schedule for the next four sessions. The VI-60 reinforcement schedule was utilized during subsequent fear discrimination and was completely independent of auditory cue or foot shock presentation.

### Pre-exposure

Each rat was pre-exposed to the three auditory cues to be used in fear discrimination in two sessions. Auditory cues were 10-s in duration and consisted of repeating motifs of a broadband click, phaser, or trumpet. Previous studies have found these stimuli to be equally salient, yet highly discriminable. Stimuli available here: http://mcdannaldlab.org/resources/ardbark. The 42-min pre-exposure sessions consisted of four presentations of each cue (12 total presentations) with a mean inter-trial interval (ITI) of 3 min and at least 5 minutes of initial habituation. The order of trial type presentation was randomly determined by the behavioural program and differed for each rat during each session.

### Fear Discrimination

For the next sixteen sessions, all rats underwent Pavlovian fear discrimination. Each 54-min session began with a five minute warm-up period during which time no cues or foot shock were presented. During fear discrimination, each auditory cue predicted a unique foot shock (0.5 mA, 0.5 s) probability: danger, *p*=1.00; uncertainty, *p*=0.25; and safety, *p*=0.00. The foot shock was administered one or two seconds following the termination of the cue on danger and uncertainty-shock trials. A single session consisted of 4 danger, 2 uncertainty-shock, 6 uncertainty-no shock, and 4 safety trials with a mean inter-trial interval of 3 min. The order of trial presentation was randomly determined by the behavioural program and differed for each rat, every session. The physical identities of the auditory cues were counterbalanced across individuals.

### Analysis

Timestamps for nose poke, cue onset, and shock onset were collected with Med Associates software. Nose poke rates (Pokes/min) were calculated in 1s bins aligned to cue onset and shock offset. One set of analyses focused on second by second nose poke rates around cue onset and shock offset. A second set of analyses focused on nose poke rates during four *a priori* periods of interest (fig 1A): baseline - 10s prior to cue onset, cue - 10s cue period, immediate post shock - first 2s after shock offset, and delay post shock - 4s period starting 3s after shock offset. Differential nose poke rates between trial-types, and trial-type elevations in nose poking over baseline were examined using 95% bootstrap confidence intervals.

**Figure 1.**
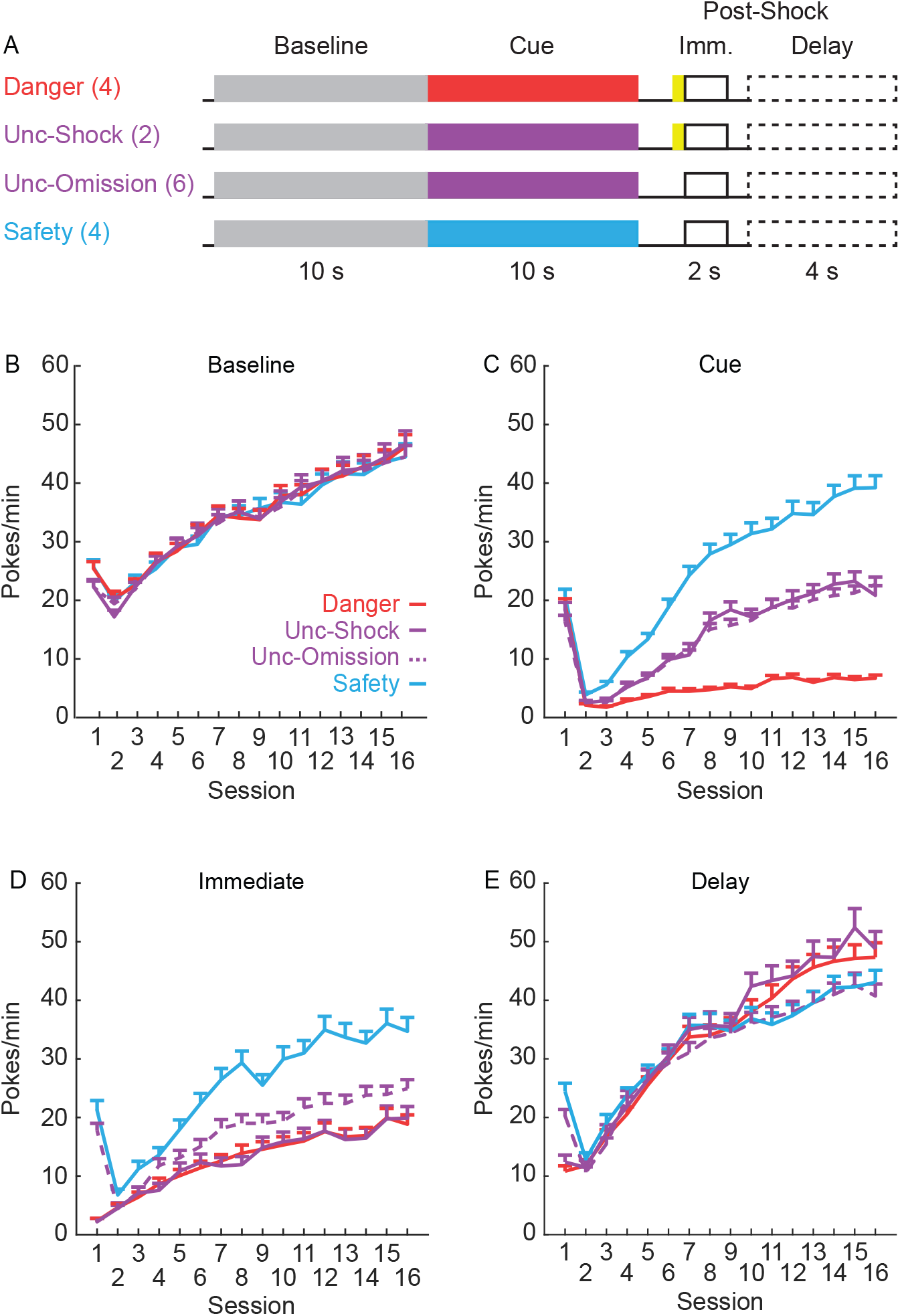
Trial schematic showing the four periods of interest for each trial-type (A) and mean rates of responding in each session of fear discrimination during baseline (B), cue (C), immediate post shock (D), and delay post shock (E) periods. Error bars show ^+^SEM.

## Results

Our laboratory has observed robust danger, uncertainty, and safety discrimination using suppression ratios calcluated from rates of rewarded nose poking^31,32,35–37^. Classic studies examining defensive behaviour have reported robust post shock freezing^18,25,38^. To determine if cue-elicited and shock-elicited behaviours are observed in a common measure, we first analyze nose poke rates for our four *a priori* periods of interest over the 16 discrimination sessions. ANOVA with factors of trial-type (danger, uncertainty-shock, uncertainty-omission and safety) and session (1-16) are conducted on the mean rates of responding (averaging across all the time bins for each period, for each subject). As expected, baseline nose poke rates do not differ between trial types. All trial types show the same gradual increase in responding over sessions (fig 1B), confirmed by a significant effect of session (F_15,1830_ = 119.77, p=1.65 ×10^−258^, η_p_^2^ = .46, op=1.0), and no significant effect of trial type (F_3,366_ = 0.80, *p*=0.49, η_p_^2^ = .007, op=0.22), or interaction between trial type and session (F_45,5490_ = 1.16, *p*=0.21, η_p_^2^ = .009, op=0.99). These results reduce concerns that trial-type differences in nose poking during cue and post shock periods are the result of differences in baseline nose poking.

Consistent with our previous results using suppression ratios, measuring rates of nose poking reveals complete discrimination of danger, uncertainty and safey^31,32,39^. Nose poking is reduced to all cues between the first and second sessions, but the emergence of fear discrimination is clearly seen thereafter. Nose poke rates show the greatest increase over sessions during safety, an intermediate increase during uncertainty and the least increase during danger (fig 1C). This pattern is confirmed by significant effects of trial-type (F_3,366_ = 234.70, *p*=6.71 × 10^−85^, η_p_^2^ = .66, op=1.0), session (F_15,1830_ = 141.16, *p*=5.73 × 10^−292^, η_p_^2^ = .54, op=1.0), and importantly a significant trial-type × session interaction (F_45,5490_ = 40.08, *p*=7.56 × 10^−299^, η_p_^2^ = .25, op=1.0).

Consistent with post shock freezing reports, nose poking is greatly reduced immediately following foot shock delivery on danger and uncertainty-shock trials during the first session. Reduced nose poking generalizes during the immediate period on all trial-types by the second session. Responding increases in subsequent sessions, with the greatest increase following safety, lesser increases following uncertainty-omission and the least increase following foot shock trial-types (danger cue and uncertainty shock, fig 1D). Again this pattern is confirmed by ANOVA revealing significant effects of trial-type (F_3,366_ = 55.03, *p*=2.21 × 10^−29^, η_p_^2^ = 0.31, op=1), session (F_15,1830_ = 77.81, *p*=1.97 × 10 ^− 183^, η_p_^2^ = 0.39, op=1.0), and a significant trial-type × session interaction (F_45,5490_ = 4.12, *p*=1.35 × 10^−18^, η_p_^2^ = .033, op=1.0).

The baseline, cue, and immediate post shock results confirm that well-known behavioural consequences of shock-associated cues and shock delivery are observed when measuring nose poke rates. A distinct pattern emerges in the delay, post shock period. Like for the immediate period, nose poking is strongly reduced between the first and second sessions. Through session 8, there is little difference in nose poke rates between the four trial-types. However, in all remaining sessions, responding *increases* following foot shock (danger and uncertainty-shock trial-types) compared to no foot shock (safety and uncertainty-omission trial-types). Foot shock facilitates rewarded nose poking during the delay period. Descriptions are confirmed by ANOVA which found significant effects of trial-type (F_3,366_ = 8.58, *p*=0.000016, η_p_^2^ = 0.066, op=0.99), session (F_15,1830_ = 136.65, *p*=3.81 × 10^−285^, η_p_^2^ = 0.53, op=1.0) and a significant trial-type × session interaction (F_45,5490_ = 5.62, *p*=7.02 × 10^−30^, η_p_^2^ = 0.044, op=1.0).

### Shock facilitation of nose poking emerges over discrimination

To further investigate differential responding on shock and no-shock trial-types, difference scores are calculated for all 16 sessions, for each *a priori* period of interest. The mean rates of responding during uncertainty-omission trials are subtracted from uncertainty-shock trials (fig. 2, upper panels) and responding during safety trials are subtracted from danger trials (fig. 2, lower panels). Meaningful differences in nose poke rates between trial-types are determined by constructing 95% bootstrap confidence intervals for differential nose poke rates. Intervals that do not include zero support differential responding between trial-types for that session/period.

**Figure 2.**
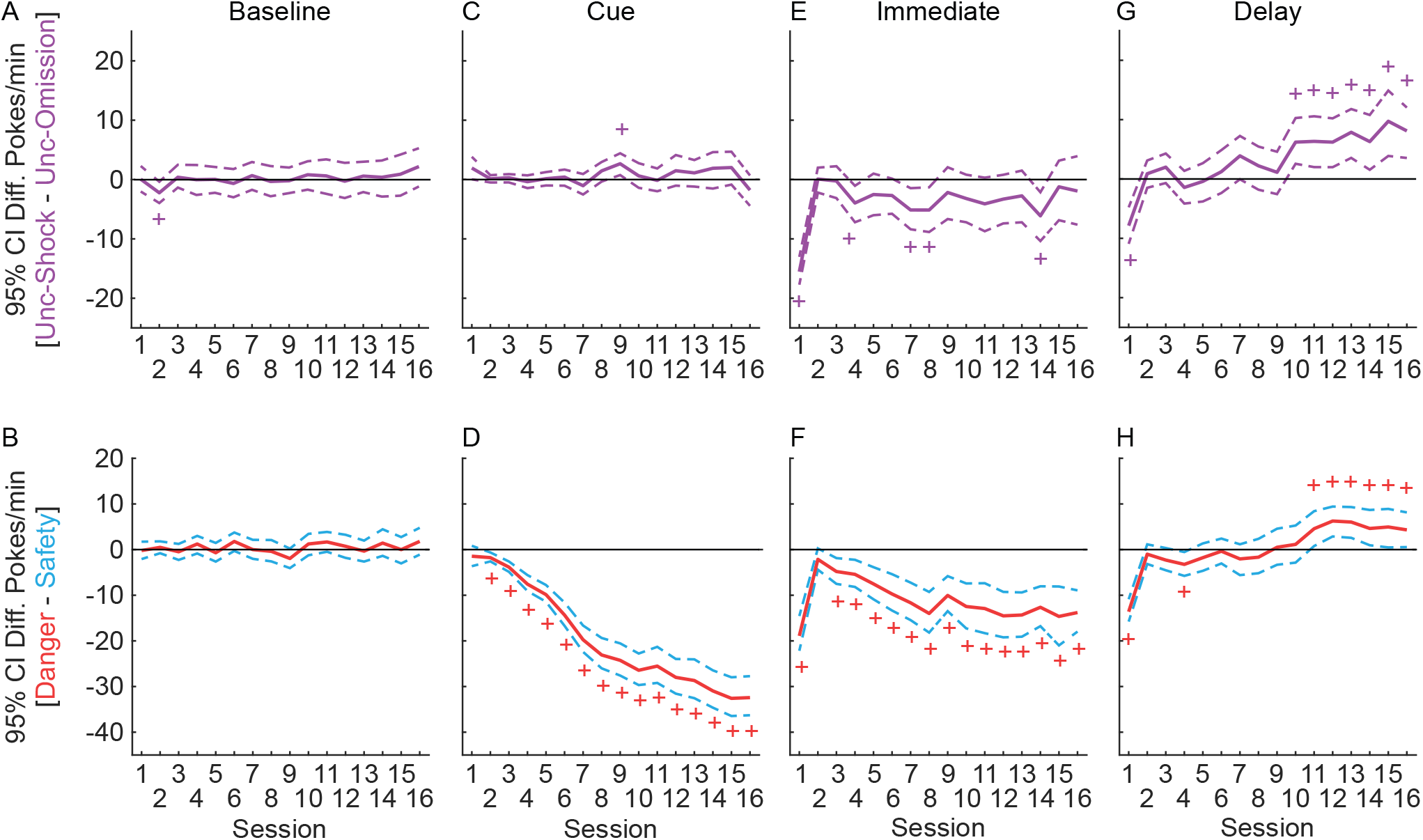
Mean difference scores and 95% bootstrap confidence intervals across sessions of fear discrimination: uncertainty-omission trials subtracted from uncertainty-shock (upper panels), safety trials subtracted from danger trials (lower panels), during baseline (A-B), cue (C-D), immediate post-shock (E-F), and delay post-shock (G-H) periods. +95% bootstrap confidence interval does not contain zero.

As expected, there is little difference in baseline responding between the two uncertainty trial types and between danger and safety trials across discrimination sessions. 95% bootstrap confidence intervals include zero (fig. 2A & B, plus marks show significance) in all but one session (unc-S - unc-O, session 2, Fig 2A, M = −2.26, 95% CI [−0.37, −3.91]). Also as expected, there is little difference between responding during the uncertainty trial-types during the cue period. Only the 95% bootstrap confidence interval for session 9 does not include zero (M = 2.64, 95% CI [4.58, 2.64]). Demonstrating rapid and robust acquisition of discrimination, rats show differential nose poke rates to danger and safety in every session except for the first (fig. 2D). The mean difference score for danger and safety increases each session, reaching a difference of −32.5 pokes/min by session 16. Note that while comparing danger/safety trial-types is an excellent indicator of discriminative cue responding, this difference will contaminate post shock responding. On the other hand, comparing uncertainty shock/omission trial-types is a poor indicator of discriminative cue responding, but is an excellent indicator of differential post shock responding.

Consistent with post shock freezing, first session nose poke rates are more greatly reduced following uncertainty shock trials compared to uncertainty omission during the immediate post shock period. Somewhat surprisingly, from session 2 onwards this difference is reduced. Differential nose poking during the immediate period hovers just below zero, with only sessions 1,4, 7, 8, & 14 showing lesser responding on shock trials (fig. 2E). Comparing danger and safety responding reveals a more consistent pattern. Responding is lower immediately following shock during danger trials, compared to the immediate period during safety trials. Apart from session 2, the 95% confidence interval does not include zero (fig. 2F). Though again, reduced nose poking immediately following danger shock presentation may have carried over from the cue period.

We are most interested in the pattern of differential responding during the delay period (fig 2G & H). For both trial-type comparisons, a considerable reduction in nose poking is observed during the foot shock delay period in the first session (95% bootstrap confidence intervals do not include zero for unc-shock vs. unc-omission and danger vs. safety). With the exception of the danger-safety comparison in session 4, no subsequent foot shock reduction in nose poking is observed. By the end of discrimination, higher nose poke rates are observed following foot shock, compared to no-shock trials. Shock delivery facilitates nose poking during the delay period. Facilitation following uncertainty-shock delivery is observed during each of the final 7 sessions (10-16), and facilitation following danger-shock delivery is observed during each of the final 6 sessions (11-16; 95% bootstrap confidence intervals do not include zero).

### Reduction, then facilitation of nose poking following foot shock

To examine the temporal emergence of nose poke facilitation following shock delivery, we focus on responding in the final six sessions. We calculate mean nose poke rates for the 2, 1-s baseline intervals, 10, 1-s cue intervals and 10, 1-s post shock intervals. As expected, no differences in baseline nose poking are observed between trial-types (fig 3A). Responding is reduced for all cues in the first interval, although differential responding is evident (safety > uncertainty-omission = uncertainty-shock > danger). From the second interval on, the full fear discrimination pattern is clear (fig 3A). ANOVA with trial-type and time (1-s interval) as factors confirm a significant effect of trial-type (F_3,366_ = 228.87, *p*=1.37 × 10^−83^, η_p_^2^ = 0.65, op=1.0), time(s) (F_9,1098_ = 15.10, *p*=2.42 × 10^−23^, η_p_^2^ = 0.110, op=1.0), as well as a significant interaction between trial-type and time (F_27,3294_ = 14.23, *p*=7.32 × 10^−61^, η_p_^2^ = 0.10, op= 1.0). Importantly, nose poke rates differ between danger and safety for every cue interval (95% bootstrap intervals do not include zero), but do not differ between uncertainty-shock and uncertainty-omission for any cue interval (95% bootstrap intervals include zero).

**Figure 3.**
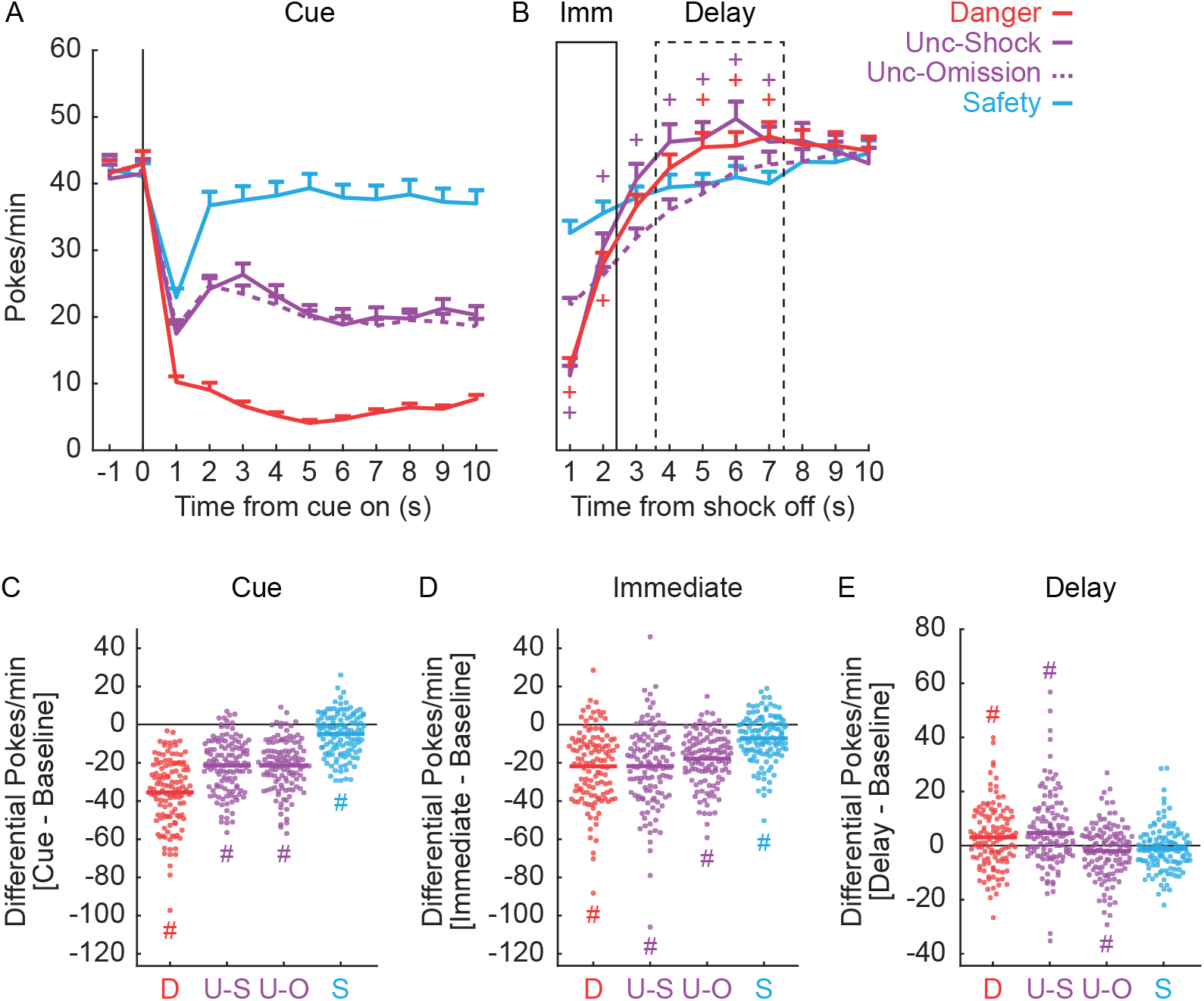
Mean rates of responding (pokes/min) in sessions 11-16 per second of: cue presentation and 2 s baseline preceding cue (A), 10 s post-shock including immediate and delay post shock periods (B). Baseline subtracted mean rates of responding for each subject in sessions 11-16 for each trial type (D = danger, U-S = uncertainty shock, U-O = uncertainty omission, S = safety), during cue (C), immediate post shock (D), and immediate delay (E) periods. ^#^95% bootstrap confidence interval does not contain zero.

The pattern of responding shows a dramatic but consistent change during the 10 s following shock offset (fig 3B). In the first interval, responding is most greatly reduced following uncertainty-shock and danger, somewhat less to uncertainty-omission and least to safety (safety > uncertainty-omission > uncertainty-shock = danger). This pattern erodes in the second interval, and responding sharply increases for uncertainty-shock and danger trials. This trend continues, with uncertainty-shock and danger nose poke rates eclipsing uncertainty-omission and safety rates, peaking ~5-6 seconds following shock offset (uncertainty-shock = danger > uncertainty-omission = safety). Nose poke rates normalize in the final ~3 seconds. ANOVA confirms a significant effect of trial-type (F_3,366_ = 6.92, *p*=0.00015, η_p_^2^ = 0.054, op=0.98), time (F_9,1098_ = 125.05, *p*=1.86 × 10^−161^, η_p_^2^ = 0.51, op=1.0), and trial-type × time interaction (F_27,3294_ = 16.40, *p*=1.64 × 10^−71^, η_p_^2^ = 0.12, op=1.0).

To reveal the differential response pattern driving the ANOVA interaction, we construct 95% bootstrap confidence intervals for differential responding during each interval: uncertainty-shock vs. uncertainty-omission and danger vs. safety (fig 3B, plus marks indicate differential nose poke rates). Confidence intervals confirm lower nose poke rates during the first post shock interval on uncertainty-shock trials compared to uncertainty-omission. Impressive to us, responding during uncertainty-shock trials exceeds that for uncertainty-omission trials in each of the next six intervals, as confirmed by the 95% confidence intervals not including zero for any of the time points. Responding during shock and no shock trial-types are equivalent during the final 3 intervals, with all the confidence intervals including zero. Comparing danger and safety trials reveal a similar pattern but with fewer intervals demonstrating facilitation. This is likely because safety responding remains high throughout the post shock period. Nevertheless, reduced responding to danger during the first two intervals gives way to enhanced responding to danger during intervals 5-7. No bootstrap confidence intervals include zero for these time points. Responding converges in the final 3 intervals, again confirmed by no bootstrap confidence interval including zero.

Observing increased responding following shock delivery, compared to trials on which shocks do not occur, provides compelling evidence of a relative foot shock facilitation effect. To determine if absolute facilitation occurs, we compare nose poke rates during each period of interest to baseline. Difference scores (period - baseline) are calculated for each trial-type/period and 95% bootstrap confidence intervals constructed fig 3C - E). Decreases in responding under baseline are observed for each trial type during the 10-s cue period (fig 3C). Similarly, decreases in responding under baseline are observed for each trial type during the immediate shock period (fig 3D). Revealing absolute facilitation, increases in responding over baseline during the delay period are observed for danger (~110% of baseline responding, M = 3.06, 95% CI [0.96, 4.96]) and uncertainty-shock trials (~115% of baseline responding, M = 4.73, 95% CI [2.03, 7.26]). Responding decreases are observed to uncertainty-omission (~99% of baseline responding, M = −1.98, 95% CI [−3.64, −0.24]), while safety responding does not differ from baseline (~102% of baseline responding, M = −1.22, 95% CI [−2.60, 0.05]).

### Shock facilitation of nose poking is a distinct behavioural mechanism

We are curious if foot shock faciliation of responding is related to other aspects of behaviour and fear discrimination. We first examine baseline nose poke, the idea being that rats showing higher baseline poking may show stronger facilitation following foot shock. To determine this, we compare sessions 11-16 mean baseline subtracted pokes during the delay period for both uncertainty-shock and danger trials (data from Fig 3E) to baseline nose poke rates from the appropriate trial-type. (Fig 4 A,B). Baseline nose poke rate is unrelated to the magnitude of foot shock facilitation. Zero relationships are observed for uncertainty-shock (R^2^ = 1.49 × 10 ^−4^, *p*=0.89) and danger (R^2^ = 0.001, *p*=0.70). Perhaps rats showing better discrimination show superior facilitation. Baseline subtracted pokes during delay are now compared to differential poke rates for danger and safety from the cue period (Fig 4 C,D). Zero relationships are observed for uncertainty-shock (R^2^ = 0.01, *p*=0.25) and danger (R^2^ = 0.01, *p*=0.25). Finally, we ask if the nose poke reduction observed during the immediate period predicts facilitation during the delay period. There may be a sort of slingshot effect, in which more strongly suppressed responding during the immediate period results in a greater ‘acceleration’ of responding during the delay period. Again, zero relationships are observed between immediate suppression and delay facilitation for uncertainty-shock (R^2^ = 0.02, *p*=0.12) and danger (R^2^ = 0.01, *p*=0.22). The only relationship we could find is within facilitation itself. Post shock facilitation during the delay period is positively correlated for danger and uncertainty-shock trials (R^2^ = 0.35, *p*=3.67 × 10 ^−13^). The results suggest that foot shock facilitation of reward seeking is a behavioural mechanism that is distinct from baseline rate of reward seeking, fear discirmination, and foot shock suppression of responding.

**Figure 4.**
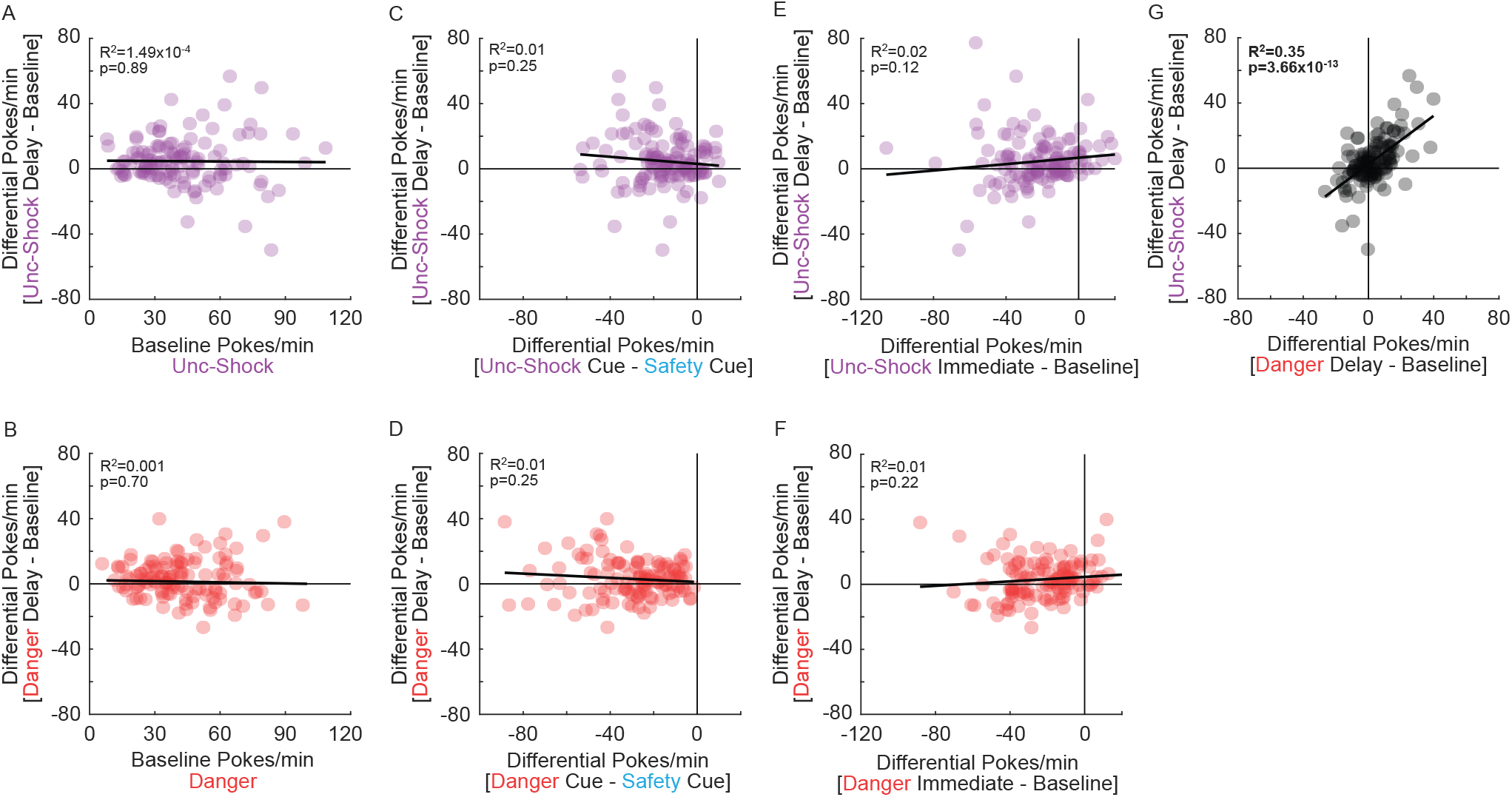
Post-shock facilitation correlated against other aspects of behaviour. Mean differential rates of responding for each subject during uncertainty shock trials (uncertainty-shock minus baseline, upper panels), and danger trials (danger minus baseline, lower panels), correlated against the baseline period (A-B), danger safety discrimination (danger minus safety, C-D), and immediate post-shock period (immediate post-shock minus baseline, E-F). Post-shock facilitation effect in the post-shock delay period correlated against other aspects of behaviour. Mean differential rates of responding for each subject during uncertainty shock trials (uncertainty-shock minus baseline, upper panels), and danger trials (danger minus baseline, lower panels), correlated against (G).

## Discussion

We set out to reveal how reward seeking around foot shock and associated cues changes as a function of experience. Measuring nose poke rate during cue presentation reveals a behavioural pattern our laboratory has previously observed with suppression ratios^31,33^. Reduced responding to all cues in early sessions gives way to discriminative cue responding: danger < uncertainty < safety. Consistent with post shock freezing^25^, early sessions are dominated by foot shock suppression of reward seeking that begins immediately following shock offset and persists thereafter. Post shock suppression diminishes as fear discrimination continues, and is mostly confined to the first two seconds immediately following shock offset. Consistent with a prior report^19^, foot shock facilitation of reward seeking emerges during later sessions. Facilitation is observed when foot shock is fully predicted on danger trials and is surprisingly delivered on uncertainty trials. Facilitation is rapid and transient, appearing as quickly as one second following shock offset and augmenting reward seeking for ~6 seconds. Foot shock facilitation of reward seeking is a distinct behavioural menchanim, unrelated to the rate of baseline nose poking, degree of discrimination and degree of foot shock suppression of reward seeking.

Before discussing theoretical accounts and practical implications, several limitations must be raised. First, both the original demonstration of facilitation and our results used male rats. The obvious question is if foot-shock facilitation of reward seeking is observed in female rats. This work is underway in our laboratory and we speculate that female rats show foot shock facilitation of reward seeking. This is based on our observation that female rats readily acquire fear discrimination^37^. Further, manipulating shock-related prediction error activity - although a distinct behavioural mechanism from facilitation - has an equivalent effect on male and female fear behaviour^34^. We speculate that facilitation - like fear discrimination and prediction error - is a behavioural mechanism conserved across sexes. Second, our results demonstrate that foot shock facilitation of reward seeking is observed within our fear discrimination procedure. However, the full conditions that produce facilitation remain unknown. It is possible that we happened to select the exact parameters (cue duration, shock duration, shock intensity, inter-trial interval, etc.) to maximize the facilitation effect. In this case our results would not generalize to other conditioned suppression procedures. Conversely, it is possible that we observed facilitation despite selecting non-optimal parameters. In this case, our present results would underestimate the magnitude of the effect. Careful, parametric studies are required to reveal the conditions that produce foot shock facilitation of reward seeking.

Various theoretical accounts have been offered to explain the increase in appetitive responding following an aversive event. A compensatory account suggests that foot shock facilitation of reward seeking occurs as a result of subjects compensating for rewards ‘lost’ during previous suppression^20^. This is a plausible explanation for facilitated responding during danger versus safety trials. Responding is suppressed throughout the danger cue, but not during the safety cue. Thus, greater responding should be observed following shock offset on danger trials to compensate for ‘lost’ rewards. The compensatory account seems unable to explain facilitation of responding following foot shock on uncertainty trials. Responding is equally suppressed during the cue period for uncertainty-shock and uncertainty-omission trial types, yet facilitation is only observed during uncertainty-shock trials.

Another possible account is learned safety, in which a cue predicts the absence or omission of an aversive event^20,40–42^. It is plausible that in the current study, shock-offset and/or the post-shock time period became associated with the absence of shock, reducing fear and suppression. However, it seems unclear how this would account for the transient period of facilitated reward seeking observed (~6s), when the actual shock-free period signaled by shock-offset is far greater (2.5-3.5 minutes). Learned safety might anticipate a much longer time period of heightened responding, that decreased as the likelihood of the next foot shock increased. Indeed, this pattern can be observed when unsignalled foot shocks are presented in regular intervals over a baseline of reward seeking^30^.

Alternatively, the opponent process theory of acquired motivation can also provide an account of post shock increases in appetitive behaviour^19,43–45^. In opponent process theory, presentation of an aversive stimulus elicits a negative hedonic ‘a process’ and is followed by an opponent positive ‘b process’. The hedonic state at a given time is the difference between the size of the a process and b process (a - b). Critically, the a process rapidly engages, peaks, and decays following shock termination. The b process is initially weaker, engaged with a delay (relative to the a process), and is slower to peak and decay. So with limited experience, the a process will dominate. Continued experience with the aversive stimulus selectively and non-associatively strengthens the b process. This strengthening permits the b process to outcompete the a process, particularly from its peak to decay. Strengthening of the hedonically positive b process with continued experience can then explain the transition from foot shock suppression to facilitation of reward seeking. Opponent process theory can also account for the transient nature of facilitation. Even though it increases in strength, the b process will still decay and terminate shortly after it is engaged. As the b process decays, reward seeking returns to baseline levels. Opponent process theory provides a reasonable explanation of the emergence of facilitation over discrimination and the time period during which facilitation is observed following foot shock.

An opponent process account lends itself well to relief learning^46–52^. Relief learning is commonly obtained through backwards conditioning, in which foot shock predicts a cue with a 1-3 s delay. Relief learning can endow a cue with inhibitory properties. For example, a backward conditioned cue can diminish startle, while a forward conditioned cue can potentiate startle^49^. The ‘event’ supporting inhibitory learning in backward conditioning is thought to be the transient relief generated by the cessation of the painful stimulus. This phenomenological description of relief is strikingly similar to the positive hedonic state elicited by the b process in opponent process theory. Although speculative, our results suggest that the positive, post shock signal that supports relief learning is sufficient to facilitate reward seeking on its own. Foot shock facilitation of reward seeking may be a form of general affective Pavlovian to instrumental transfer^53,54^ that bridges appetitive and aversive motivational systems^55^.

From a practical perspective, the transition from foot shock suppression to facilitation may be adaptive, capitalizing on the subject’s knowledge of the environment. When an aversive event is first experienced, it is impossible to predict when subsequent events will occur, much less the nature of those occurrences (number, duration, intensity, etc.). Engaging neurobehavioural systems for threat and negative affect is adaptive, minimizing detection and further risk of harm. Continued experience may allow the subject to predict not just when an aversive event will occur, but the specific nature of each occurence. Now it is adaptive for termination of the aversive event to engage neurobehavioural systems for positive affect, facilitating reward seeking.

Associative learning has been a valuable tool in identifying adaptive behaviour that becomes maladaptive in stress and anxiety disorders. Fear discrimination and safety learning procedures have revealed altered behavioural responding to threat and safety cues in post-traumatic stress disorder^56–58^. Our results and extant findings strongly suggest that extending associative learning analyses to responding around aversive events will be equally valuable^59,60^. Preclinical research can identify brain regions permitting foot shock facilitation. Relief learning studies have already identified critical roles for the ventral tegmental area and nucleus accumbens shell^48,49,61,62^. The mesolimbic dopamine system is perhaps likely to contribute to foot shock facilitation of reward seeking. Clinical studies can reveal experience-dependent changes in behaviour around aversive events. A straightforward hypothesis is that the suppression → facilitation transition normally observed for aversive events is slowed in individuals with stress and anxiety disorders. Impairment could result from a failure to transition from aversive processing in nociceptive regions^63^ to reward regions^46,64^. Of course, many more outcomes are possible. Revealing the neurobehavioural mechanisms underlying foot shock facilitation may allow us to harness the positive power of aversive events to promote adaptive behaviour.

## Acknowledgements

The authors declare no conflicts of interest.

